# Mixing is required for uniform reconstitution of filter-dried protein antigens in a single-injection vaccine formulation

**DOI:** 10.1101/247403

**Authors:** Napawan Thangsupanimitchai, Alexander D. Edwards

## Abstract

Ambient temperature filter dried vaccine formulations have been proposed to simultaneously achieve thermostability and offer a ready-to-use immunisation device that combines reconstitution and injection. Vaccine concentration should be uniform at the point of injection, but the uniformity following direct reconstitution of filter-dried vaccines has not been reported. We present here a study of vaccine mixing and release following dissolution of filter-dried model protein and toxoid antigens within a single syringe, filter and needle unit. Release was better for filters made from glass than cellulose. Without additional mixing, uniformity was poor and only 41% of input protein was released from protein filter-dried onto glass fibre. In contrast, adding a simple glass bead and mixing by inversion, 100% release antigen solution was achieved, with uniform concentration at exit from the needle throughout a simulated injection. Adsorption onto alum adjuvant had no detectable effect on vaccine dissolution and mixing. The uniformity and yield of low doses of diphtheria and tetanus toxoid was also improved by mixing, albeit with a lower yield of 60-68%. We conclude that uniformity and mixing should be studied to ensure safety and efficacy of directly reconstituted filter-dried vaccine formulations.

## 1 Introduction

A significant proportion of the cost of vaccination programmes lies in safe refrigerated storage and distribution, and costs of trained healthcare professionals for administration. This has driven extensive research into drying and thermostabilisation technology for vaccines [1]. Traditionally freeze-dried vaccines still require refrigerated storage, although excipient optimization can achieve thermostable lyophilized vaccines [2]. Freeze-dried vaccines still require careful reconstitution and preparation prior to administration, typically reconstitution in a freeze-drying vial, followed by transfer to a syringe and needle for injection. Whilst the widespread use of liquid formulations distributed in pre-filled syringes for many common vaccines reduces staff time and skill level, liquid vaccine preparations are inevitably less stable than freeze-dried formulations, with solid formulations more stable and easier to handle [3, 4] because chemical (e.g. hydrolysis) and physical (e.g. foaming or aggregation) instability have far faster kinetics in aqueous form. Pre-filled syringes could be argued to be more expensive formulations than traditional lyophilized vials, but in reality freeze-drying is already a labour intensive and slow process constrained by capacity of industrial freeze-driers [3], and the processes used for manufacture of vaccines in pre-filled syringes are now well developed. Furthermore, lyophilized vaccine vials still require water for injection plus disposable syringe and needle, which whilst low-cost components, increase the complexity of the supply chain. The ideal vaccine formulation must therefore be not only produced from low cost components, but must also have cost-effective scalable manufacturing process, and the simplest possible format for the end-user administrating the vaccine dose. To give the product stability and avoid cold-chain storage, this device would contain a dry physical form produced using a scalable drying method which maximises thermostability. Whilst alternatives to injection such as dried oral solid dosage forms [5] offer benefits, the overwhelming majority of current vaccines require injection. Reconstitution of dried vaccine to a liquid for injection therefore becomes an important engineering consideration.

The cost of freeze-drying, and burden of cold-chain storage and complex preparation for administration has driven research into a range of novel formulations, and modelling has shown that thermostable vaccines would deliver significant improvements to the vaccine supply chain[6]. Multiple alternatives to freeze-drying have been explored, such as spray- or foam- drying [7]. A long-established and intensively studied example is the use of ambient temperature drying to form an amorphous sugar glass from high glass transition temperature carbohydrates such as trehalose [8–10]. This method has been combined with simplified formulation to deliver a thermostable, single-component vaccine formulation and injection format that shows great promise [11, 12]. The vaccine is dried to form a sugar glass on a non-woven fibre filter substrate, which is then encased in a filter housing which is connected between a syringe filled with water for injection, and the needle [13, 14]. The sugar glass vaccine dissolves in flow as the syringe is depressed, and the dissolved vaccine injected directly into the patient. This approach is suitable to fragile live attenuated recombinant viral vaccines including modified vaccinia and adenovirus, which were preserved on the membranes for at least 6 months storage at high temperature (45°C) [13]. Similarly, hepatitis B vaccine dried onto filters are stable and immunogenic at 55°C for 7 weeks [9].

Tetanus and diphtheria toxins derived from *Clostridium tetani* and *Corynebacterium diphtheriae*, respectively are chemically inactivated to form toxoids that retain antigenic structure and potently induce a protective toxin-neutralizing immunity. Whilst very stable when compared to live attenuated whole viral vaccines, structure can still be lost on storage or during processing or formulation [15, 16] Toxoids such as tetanus toxoid are among the most heat-stable biological vaccine components. However, World Health Organisation (WHO) recommend tetanus and diphtheria toxoids still require storage at 2-8°C for years, with reduced stability of months at 25°C and only weeks at 37°C [15]. Therefore, these toxoids were selected as a suitable model for study of reconstitution from filter-dried vaccine formulations.

As this highly promising “all-in-one” formulation is now being developed for use in new human vaccine products, we questioned if the dissolution characteristics of reconstitution from the filter has been studied in enough detail. The uniformity of any drug prior to injection into patients is critical to safe and controlled administration and function, and we wished to understand if this novel formulation can deliver a uniform concentration of reconstituted vaccine. Whilst fluid flow through the filter matrix will generate a degree of turbidity, and some mixing will occur between reconstitution from the filter and the end of the needle, there is no mechanism driving axial redistribution if flow is unidirectional from syringe, through filter, into needle.

We present here a pilot study of mixing following reconstitution of filter-dried protein antigens. We used an extremely simple and low-cost injection rig to explore the feasibility and effectiveness of active mixing within a syringe prior to injection. We found that the dried vaccine is unlikely to be mixed effectively following reconstitution, and that mixing not only increased uniformity of vaccine delivered, but increased the yield. To permit not only precise quantitation of vaccine dissolution but also direct imaging of uniformity of vaccine solution, we studied fluorescently labelled protein and alum-adsorbed protein and diptheria and tetanus toxoid as model vaccine antigens.

## 2 Materials and Methods

### 2.1 Materials

To make fluorescent model protein antigen, BSA at >=2 mg/ml in 0.5 M of carbonate buffer (pH 9.5) and 10 mg/ml FITC in anhydrous dimethyl sulfoxide (DMSO) were combined a ratio of 80-160 µg FITC per mg of antibody. After incubation at room temperature for 1 hour, conjugates were purified by size-exclusion chromatography. BSA, FITC and Sephadex G-50 superfine were from Sigma-Aldrich (Dorset UK). Phosphate buffer saline tablets (Sigma, Dorset UK) were dissolved in ultrapure water and ProClin 300 (Sigma, Dorset UK) was added as a preservative. Adjuvant adsorbed antigen was prepared following manufacturers direction, briefly Imject Alum (Thermo Fisher UK, Paisley UK) was added to FITC-BSA at a final volume ratio of alum to immunogen of 1:3, then mixed together for 30 minutes for efficient adsorption. Non-adsorbed diphtheria and non-adsorbed tetanus toxoid were used as model antigens in a simulated combined vaccine, and were obtained from the National Institute for Biological Standards and Control (NIBSC, Potters Bar, UK) as well as monoclonal and polyclonal antibodies for sandwich ELISA antigen quantitation; all were reconstituted following the supplier’s recommended method. Two types of filter were compared for filter drying of model vaccine antigens: cellulose filters were Whatman™ qualitative filter papers grade 1 (Cat No 1001-055) with 180 µm thickness; glassfibre filters were grade 8964 with 429.3 µm thickness (Ahlstrom, Lyon, France).

### 2.2 Preparation of ambient temperature dried model antigen filters and combined injection device

A simple prototype combined injection device was made using two filter-dried vaccine antigen squares loaded into the barrel of a 2ml plastic disposable syringe (BD, Berkshire UK). To promote mixing, a 3mm diameter glass mixing bead taken from a Monovette^®^ 1.2mL blood collection tube (Sarstedt; Leicester UK) was added with the filters (Fig. 1A). To make filter-dried antigen squares, 10µL of fluorescein, fluorescent protein or antigen solution was added per 5mm square filter, and dried at ambient temperature (typically 20° C) in a petri dish until completely dry, which took a minimum of 1 hour (confirmed by kinetic analysis of mass loss). To establish methods and directly image reconstitution, 1mM sodium fluorescein solution was dried onto filters. This was followed by study of filters loaded with fluorescently labelled BSA, used as a model protein to more accurately reflect vaccine antigen dissolution. A 3.2mg/mL solution of BSA-FITC was found to be the lowest concentration suitable for quantitative imaging in the chamber. To determine if a real toxoid vaccine antigen behaved similarly to the high concentration of model protein used, both diphtheria and tetanus toxoids were diluted in ultrapure water to the lowest concentration that permitted quantitation by ELISA. A concentration of 4 lf/ml was selected for both toxoids mixed in a ratio of 1:1, and 10µL was pipetted on a 5x5 mm square of filter to yield 0.02 lf of diphtheria and tetanus toxoid per filter. Toxoid concentrations are presented in flocculation value per ml (lf/mL). When two filters in a syringe were reconstituted with 500 µl ultrapure water, the maximum concentration of reconstituted vaccine expected was 0.08 lf/mL.

**Figure 1:**
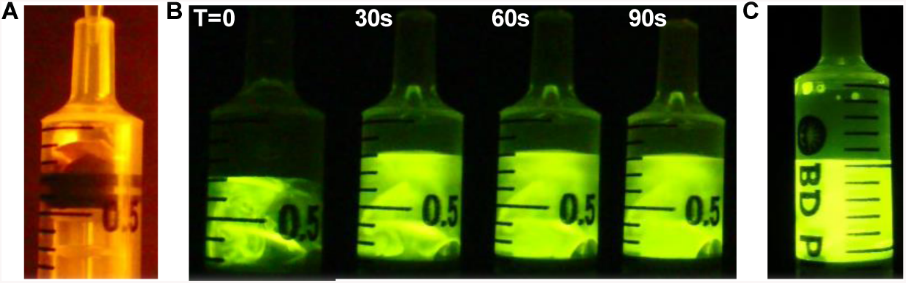
Lack of uniformity of reconstituted fluorescein solution without mixing. 10µL 1mM sodium fluorescein was dried onto filters, and placed in the barrel of a 2 ml syringe (A) and imaged as 0.5 ml of ultrapure water was added (B). The syringe was then mixed by vigorous shaking, showing uniform fluorescence when well mixed (C).

### 2.3 Measurement of mixing and dissolution of filter dried model vaccine antigens

To observe fluorescent protein mixing directly after reconstitution, filters with dried 1mM sodium fluorescein were imaged in a dark box using blue light excitation through an amber acrylic emission filter (IOrodeo, San Diego, USA) using a Canon S120 digital camera using manual aperture and exposure control. To observe fluorescent protein concentration delivered through the needle during a simulated vaccine injection, a chamber was made between a black acrylic and a clear polycarbonate sheet using two layers of pressure-sensitive adhesive tape to form a 0.2 mm gap (Fig 2A). The tape was cut to form a shape that allowed the solution to be imaged within a 20µL 2 mm high and 10 mm long rectangular space immediately after leaving the needle. Following addition of0.5 mL water with or without mixing, the syringe was attached to a 25 mm long 23 Gauge needle, bent through 90 degrees and connect to the imaging chamber. As the syringe plunger was depressed the chamber was imaged using the camera, dark box, and excitation source plus emission filter described above. The syringe barrel was imaged to define volume delivered. A dried filter with standard dose of fluorescein was included beside the chamber to permit comparison of intensity between different tests; in all tests the intensity of this standard sample was equal, confirming excitation intensity and camera sensitivity was reproducible. All videos and images were processed to evaluate uniformity and concentration of released protein using ImageJ [17].

**Figure 2:**
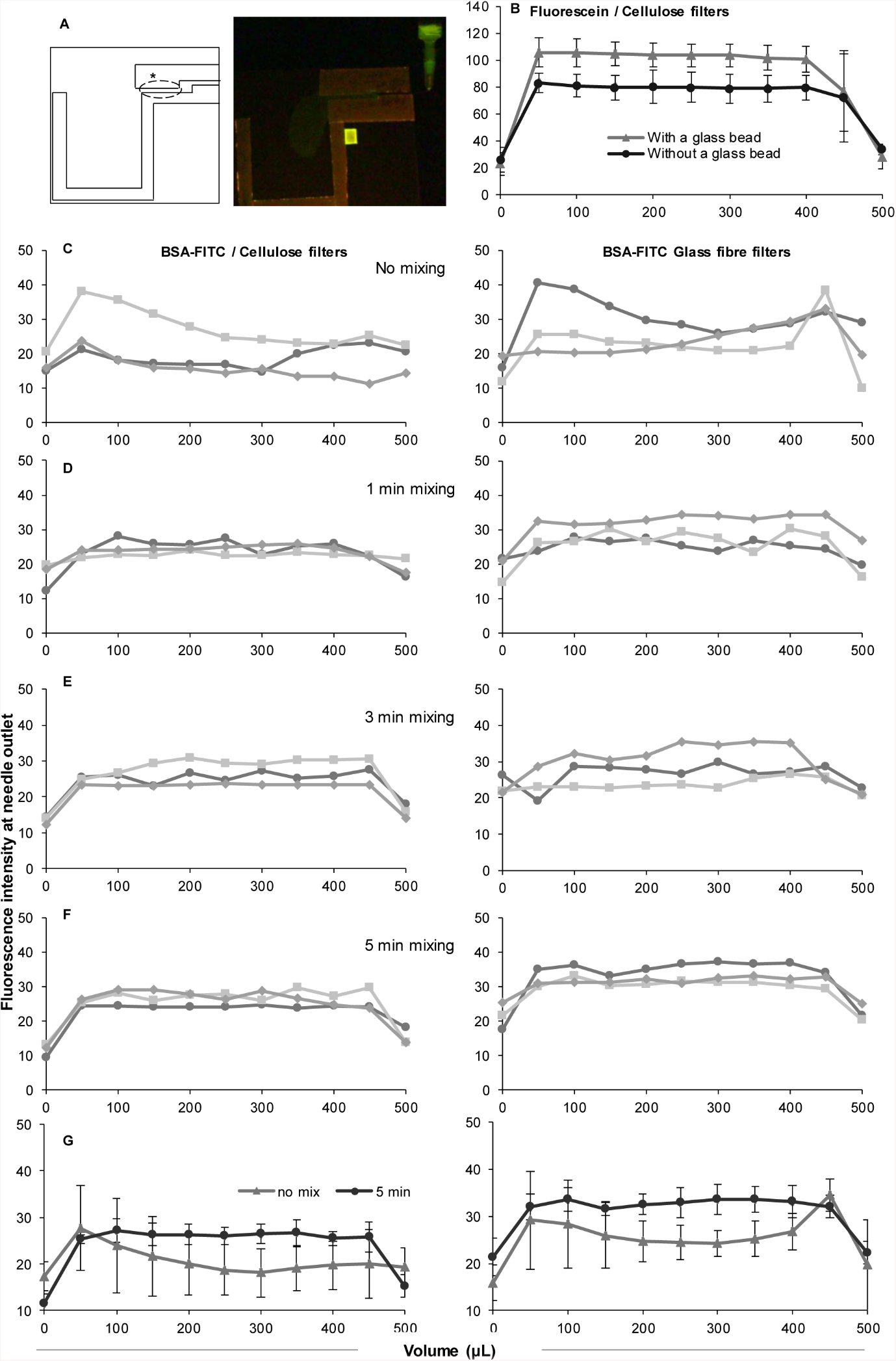
Mixing by inversion with glass bead gives uniform delivery at needle exit of model protein antigen from filter-dried model vaccine dose. **A** Diagram and example image of injection apparatus to measure protein concentration at needle exit during injection. **B** Effect of glass bead on mixing for sodium fluorescein. **C-F** Three individual replicate plots of BSA-FITC intensity at needle exit during 500µL injection from cellulose and glass filters with no mixing (C) or the indicated duration of inversion (D-F). **G** Comparison of mean and standard deviation of triplicates showing increased intensity and reduced variance with mixing vs no mixing. Error bars indicate +/− standard deviation of three replicate tests.

After the syringe was completely emptied the total yield of model antigen in the ~0.5mL solution delivered was measured using three methods. Firstly, release of BSA-FITC with or without adsorption onto alum was measured using protein determination with the Pierce 660 nm Protein Assay following manufactures instructions. Briefly, 10 µL samples were mixed with 150 µl of Pierce 660 Protein Assay Reagent (Thermo Fisher Scientific) in a microplate, and after incubation at a room temperature for 5 minutes the absorbance at 660 nm recorded and compared to a standard curve prepared from known BSA or FITC-BSA samples. When adsorbed onto alum adjuvant, protein concentrations were calculated from a standard curve of alum-adsorbed BSA. Secondly, absorbance at 495 was measured in parallel to determine fluorescein concentration; the release profiles were very similar to that observed for protein content. Thirdly, total antigen release from filter-dried toxoid squares was measured by sandwich ELISA using reported protocols for diphtheria [18] and tetanus [19] with modified blocking (2% BSA) and substrate (SigmaFast OPD; Sigma, Dorset UK). The anti-diphtheria monoclonal capture antibody was NIBSC code: 10/130, and guinea pig polyclonal detection antibody was NIBSC code: 10/128. For tetanus, monoclonal capture antibody was NIBSC code: 10/134 and guinea pigs polyclonal detection antibody was NIBSC code: 10/132. Where indicated, a two-sample students t-test was used to assess statistical significance, and statistically significant differences where *p* < 0.05 marked with *.

## 3 Results and discussion

### 3.1 A uniform flow of protein requires mixing

We initially used fluorescein dye, BSA-FITC conjugates and toxoid antigens to study the mixing and uniformity following direct reconstitution from filters in a simple reconstitution device. When imaged immediately after addition of ultrapure water to filter-dried fluorescein we found clear variations in intensity, in contrast to a bright and homogenous solution after vigorous mixing in the syringe (Fig. 1). This indicated that an additional mixing step should be incorporated to ensure vaccine is homogeneous on injection. Since mixing can occur in the syringe and needle during injection, and mixing is hard to quantify within the syringe body, an imaging chamber was made to measure fluorescence directly as the solution exits a needle suitable for vaccine administration (Fig. 2A). Concentration was quantified within a uniform rectangular 20 µL chamber that was 2 mm high, 10 mm long, with a thickness of 0.2 mm. Glass beads are used to mix anticoagulant into small volume blood samples in products such as Monovette®. In pilot tests, the effectiveness of adding a glass mixing bead and mixing by inversion was assessed with filter-dried fluorescein dye. Inversion mixing with a glass bead gave a higher average fluorescence intensity and smaller standard deviation than inversion without the bead (Fig 2B), and the bead was therefore included in all subsequent experiments. Pilots also confirmed that dissolution from filters differs somewhat between the small molecule dye fluorescein vs protein, and so subsequent experiments used fluorescently labelled model protein.

BSA-FITC was chosen as a model protein antigen and filter-dried onto cellulose and glass fibre squares. The release profile was assessed by measuring intensity of sequential 50µL volumes during injection through the chamber, after reconstituting with or without mixing. Both uniformity of intensity and overall yield were improved by mixing. Without mixing, there was significant variation both between replicates, and in the concentration delivered per unit volume during the injection period (Fig. 2C). Spikes in concentration were often detected for individual samples (for example in the first and/or last volume). This variation was reduced following mixing for any length of time, for both cellulose and glass filters (Fig. 2C-F). With glass filters, mixing also gave a clear increase in intensity. Although mixing also improved uniformity for cellulose filters, the fluorescence intensity was somewhat lower than glass filters, suggesting less efficient release. When mean and standard deviations were plotted, the reduction in variance and increase in released concentration were clear throughout the duration of the simulated injection (Fig. 2G).

### 3.2 Maximal yields are obtained from glass filters mixed by inversion with glass bead

Whilst the imaging system was ideal to determine uniformity in concentration as the injected volume passes out of the needle, and showed that variation was eliminated by mixing, it was important to determine independently the overall protein yield and determine if the apparent increase in concentration following mixing could be confirmed. The entire 500µl injection was therefore withdrawn from the reservoir, and protein yield measured. This clearly showed that the imaging system underestimated the increase in protein release following mixing, and also confirmed that the best yield was achieved by a combination of glass filters and mixing (Fig 3A). No further benefit was seen increasing mixing time above 1 minute. These observations were confirmed by absorbance measurements to quantify fluorescein concentration in bulk solutions recovered after complete injection. Likewise, when calibrated against standard concentrations imaged in the same chamber, similar total concentrations were estimated from the fluorescent imaging chamber to that found by bulk protein determination (data not shown).

**Figure 3:**
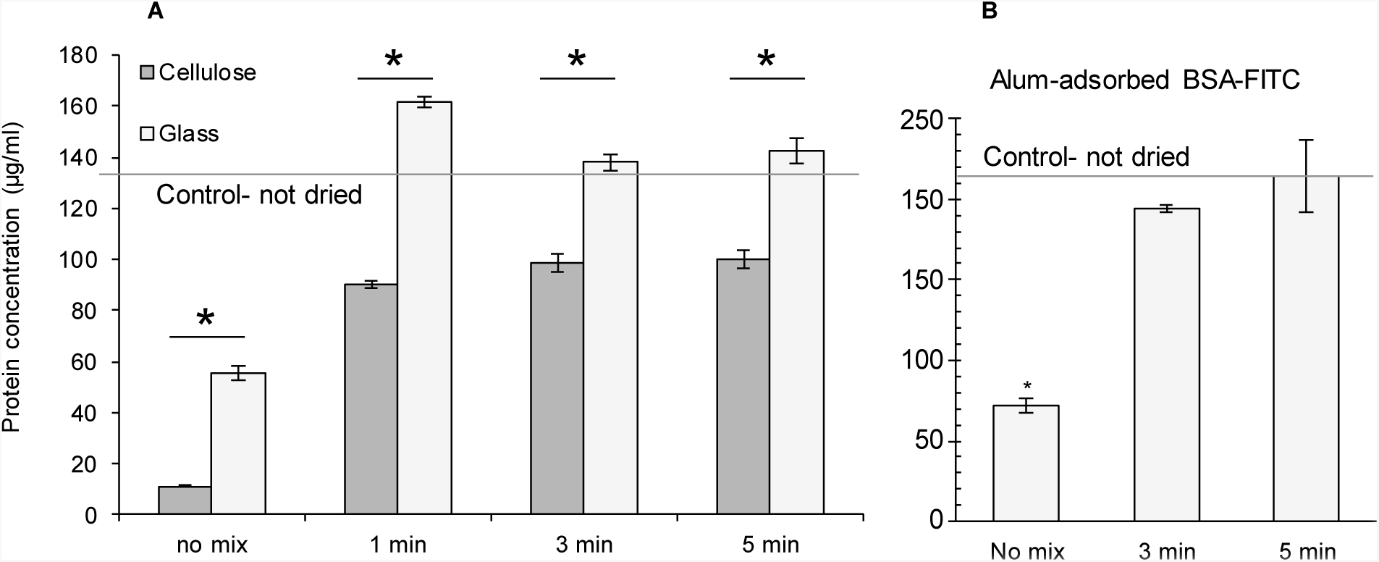
Mixing increases yield from filter-dried model protein antigen. **A** The entire 500 µL volume imaged in Fig 2 was collected for BSA-FITC samples, and antigen yield measured by protein determination. **B** BSA-FITC was adsorbed onto alum antigen and dried onto glass filters, followed by assembly into prototype single-injection devices and reconstitution with or without mixing as indicated. Total antigen yield was measured in bulk 500 µL volume by protein determination. The line indicates maximal possible recovery and is the protein concentration from a control non-dried sample of BSA-FITC diluted directly into 500 µL alongside filter reconstitution. Bars indicate mean and error bars 1 standard deviation of three independent replicates. ^*^ indicates statistically significant differences (p<0.05) calculated using a student T-test.

Protein antigens are typically weakly immunogenic without adjuvant; it was therefore important to assess the effect of the adjuvant on filter-dried protein release. After adsorbed the fluorescent protein with aluminium hydroxide, the same protein measurement procedure, was performed and once again significantly lower protein concentration was recovered without mixing. Again, a high yield of alum-adsorbed fluorescent protein was released with mixing by inversion with a glass bead (Fig 3B). Although a slightly low protein concentration was observed in the three-minute mixing group, no significant difference was found between this and control group.

### 3.4 Mixing is required for maximal recovery of low concentration filter-dried toxoid antigen

Antigen concentration for some subunit vaccines is substantially lower than that used in the imaging studies, making it far more challenging to either visualize protein release or measure protein content directly. Toxoid vaccines are potent immunogens and so very low concentrations are formulated in diphtheria and tetanus vaccines (typically around 1 lf per dose), and the reference toxoid samples we used in this study were supplied with a high concentration of protein carrier, further complicating measurement. The low concentrations plus the presence of protein carrier made it challenging to label for direct imaging or to measure antigen concentration and uniformity by fluorescence or protein determination. Instead, we exploited sandwich immunoassays developed specifically for quantitation of antigenically intact toxoids for quality control. The high analytical sensitivity of sandwich ELISA permitted us to test toxoid concentrations at or below that required for human vaccine dose. A very low-dose diphtheria and tetanus combined toxoid vaccine prototype incorporating glass fiber filter-dried antigen samples was prepared and tested and antigen content evaluated by sandwich ELISA. Significant variations in antigen content were found in the unmixed group for both diphtheria and tetanus toxoids (Fig. 4). Although an increased yield with lower sample-sample variation was observed with mixing, the yield was still substantially lower than that observed with BSA-FITC (Figs. 3 and 4). Approximately 60-70% of total input antigen content was recovered after five minutes of mixing, compared to the control sample which was treated identically but without filter drying and diluted in ultrapure water to produce a 500 µL sample. One explanation was that at this very low concentration of antigen, significant protein was irreversibly adsorbed onto the glass fiber filter during drying, which could potentially be avoided by pre-blocking or addition of blocking excipients. Likewise, adsorption of the toxoid onto alum adjuvant might overcome this challenge. Another possibility is that the drying and reconstitution process denatured part of the toxoid antigen, rendering it undetectable in the sandwich ELISA that is designed to recognize antigenically intact toxoid. However, the improved uniformity and increased yield with mixing confirmed our observations with high dose fluorescent model protein that mixing is important for uniformity and yield.

**Figure 4:**
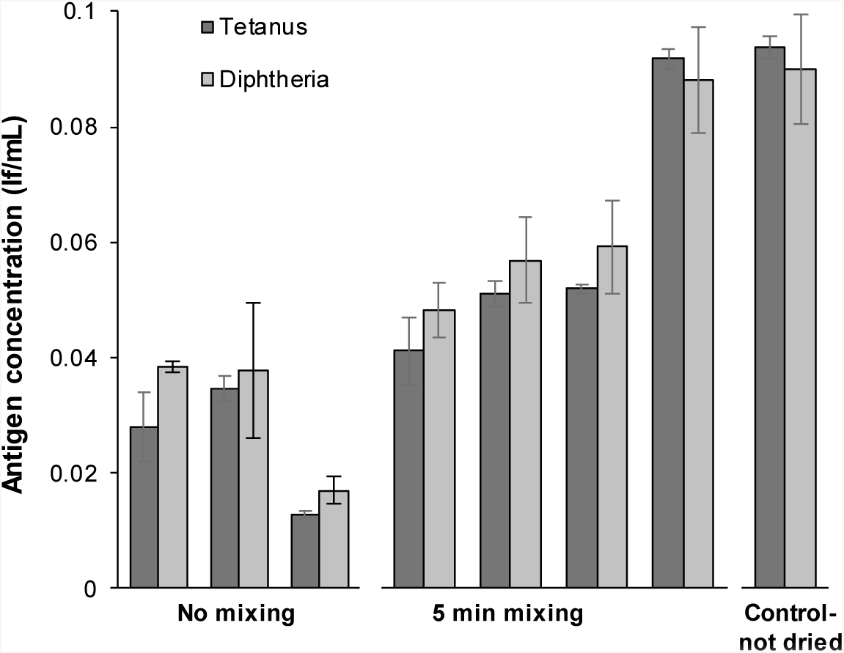
Mixing increases yield and reduces variability of release of low dose toxoid antigen, but with lower recovery. 10µL of a 2 lf/mL diptheria and tetanus toxoid mixture was dried per filter. This dose was the lowest concentration that permits quantitative measurement of release by sandwich ELISA, at this concentration a human vaccine dose would be 0.5mL. Each bar presents mean and standard deviation of triplicate ELISA measurements for one syringe containing 2 filters plus a mixing bead. The control sample is an identical filter plus 10 µL toxoid solution, which was not dried but directly diluted to 500 µL in parallel; the theoretical maximum concentration with complete release would be 0.08 lf/mL.

## 4 Conclusions

In conclusion, we suggest that direct reconstitution and injection of protein antigens from a sugar glass dried at ambient temperature onto a filter does not deliver a uniform solution, and may not be appropriate for vaccine injections. We observed fluctuations and inconsistency in fluorescence intensity in the unmixed solution both within the syringe barrel (Fig 1A) and on exiting the needle during simulated injection (Fig 2). In contrast, after even 1 minute of mixing by inversion with a glass mixing bead a decreased standard deviation of solution exiting the needle was achieved, combined with an increase in protein antigen yield (Fig 2-4). Surprisingly, adsorption onto alum adjuvant had no effect on protein reconstitution and recovery. Even with mixing, recovery of toxoids dried at very low concentrations (representative of some potent human vaccines) was lower than yields of fluorescently labelled BSA dried in microgram quantities, suggesting that each vaccine requires individual optimisation. We found a 2 ml disposable syringe can be used as a simple and low-cost housing for filter-dried vaccine, if a 3 mm glass bead is included and the device is mixed by inversion following addition of water for reconstitution. This simple single unit device would permit simpler reconstitution and distribution than conventional vial lyophilized doses. This pilot investigation suggests that reconstitution dynamics should be scrutinized for all single-injection devices intended to exploit direct reconstitution of thermostable dried vaccines.

## Acknowledgements

NT was supported by the Government Pharmaceutical Organisation (GPO), Bangkok, Thailand.

---

**Graphical Abstract.**
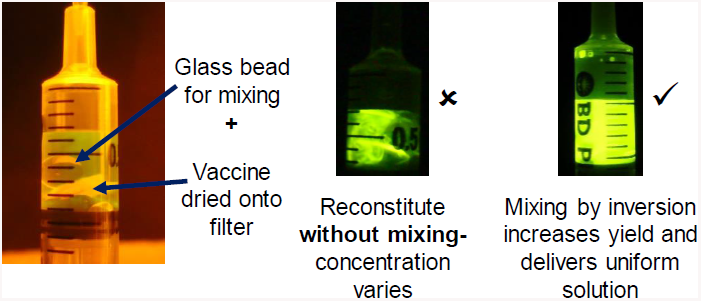

